# Accelerated Epigenetic Ageing in Major Depressive Disorder

**DOI:** 10.1101/210666

**Authors:** Heather C Whalley, Jude Gibson, Riccardo Marioni, Rosie M Walker, Toni-Kim Clarke, David M Howard, Mark J Adams, Lynsey Hall, Stewart Morris, Ian J Deary, David Porteous, 23andMe Research Team, Major Depressive Disorder Working Group of the Psychiatric Genomics Consortium, Kathryn L Evans, Andrew M McIntosh

## Abstract

**Background:** Major depressive disorder (MDD) is a severe, heritable psychiatric disorder associated with shortened lifespan and comorbidities of advancing age. It is unknown however whether MDD is associated with accelerated biological ageing relative to chronological age. This hypothesis was tested using the epigenetic clock as a measure of biological age.

**Methods:** To address the main hypothesis, using peripheral blood, we derived measures of Epigenetic Age Acceleration (EAA) in 3,833 controls and 1,219 MDD cases based on Hannum and Horvath epigenetic clocks in Generation Scotland (GS:SFHS, mean age 48 years, std dev 14.5). Models controlled for relatedness, sex, cell counts, and processing batch (basic model), as well as additional covariates of smoking and drinking status, and body mass index (BMI) (full models).

**Results:** Accelerated epigenetic ageing was found in MDD cases versus controls using the Horvath clock (β=0.0804, p=0.012 equivalent to 0.20 years) in both the basic and full models. Significant MDD*age interactions indicated greatest effects at younger age ranges. No significant differences were observed for the Hannum clock. BMI was the only additional covariate found to attenuate the relationship between EAA_Horvath_ and MDD. Further, genetic correlation analysis indicated significant overlap in the genetic aetiology of EAA_Horvath_ with BMI (r_G_=0.20, p=0.03), between MDD with BMI (r_G_=0.10, p=9.86×10^−6^), but not between EAA_Horvath_ and MDD (r_G_=0.14, p=0.125). Mediation analysis indicated partial mediation of the relationship between EAA_Horvath_ and depression status through BMI (β =0.0028; p=0.0248, ~13%).

**Conclusion:** These data imply that accelerated biological ageing is associated with MDD and partially mediated through BMI.

## Introduction

Major depressive disorder (MDD) is a heritable and disabling psychiatric condition that has recently become the leading cause of disability worldwide (1). Depression is associated with a number of age-related co-morbid physical conditions, including atherosclerosis, heart disease, hypertension and stroke (2). Further, MDD is associated with reduced life expectancy (3). Faster biological ageing in MDD has been suggested as one possible mechanism for these associations (2, 4).

Several lines of convergent evidence support the premise of accelerated biological ageing in MDD. Firstly, genetic evidence demonstrates a significant overlap between molecular pathways implicated in depression and normal ageing (2, 4–6). In some studies, particularly in the domains of memory and psychomotor speed, MDD and ageing are also both associated with reduced cognitive function as well as having overlapping structural and functional brain changes (5). Telomere shortening, a measure associated with reduced lifespan and hence proposed as an objective measure of ageing, is also found in some studies of individuals with MDD (7, 8).

DNA methylation changes are strongly associated with the ageing process and may give rise to downstream changes in gene expression and function (9). More recently, studies addressing the dynamic nature of DNA methylation with age have identified various sites within the genome where changes preferentially occur (9). DNA methylation levels at particular sites can be used to derive a score that is highly correlated with chronological age (10–12). These ‘epigenetic clocks’ have also been shown to predict all-cause mortality and lifespan (13–15); those with faster running clocks relative to chronological age have a reduced life expectancy. These peripherally-derived patterns of DNA methylation therefore provide an important instrument for the investigation of biological ageing in MDD.

We primarily sought to determine if MDD status was associated with acceleration of biological ageing. We employed two commonly used measures of DNA methylation age (DNAm age): the Hannum and Horvath clocks (10–12). We hypothesised that MDD would be associated with a higher DNAm age relative to chronological age, otherwise known as epigenetic age acceleration (EAA). We sought to test these predictions in a large population-based cohort study, the Generation Scotland: Scottish Family Health Study sample (GH:SFHS) (16, 17).

## Methods and Materials

### Study Participants

#### Generation Scotland: the Scottish Family Health Study (GS:SFHS)

GS:SFHS is a family and population-based cohort study with participants recruited at random through general medical practices across Scotland. All participants were asked to refer at least one relative to the study, but neither recruitment nor referral of a relative was dependent on the diagnostic status of any particular condition or health outcome. Initial data collection took place between 2006 and 2011. The complete study protocol is described in detail elsewhere (16–18). Brief details of assessments relevant to the current study are summarised below. Ethical approval was provided by NHS Tayside Research Ethics Committee (05/S1401/89) and written consent for the use of data was obtained from all participants.

The full GS:SFHS cohort consisted of 23,690 individuals (>18 years of age at recruitment). The present study includes 5,052 individuals on whom DNA methylation data was acquired at baseline and subsequently analysed as part of a follow-up study of GS:SFHS participants, ‘Stratifying Depression And Resilience Longitudinally’ (STRADL). Briefly, individuals were selected for DNA methylation profiling based on the availability of existing longitudinal data. Individuals who participated in the Aberdeen Children of the 1950s and Walker birth cohort studies were preferentially selected (19–21), followed by those who provided further face-to-face or questionnaire-based phenotyping of depression status.

#### Clinical assessment in GS:SFHS

MDD was diagnosed using the structured clinical interview for the Diagnostic and Statistical Manual of Mental Disorders (SCID) (22). A brief screening questionnaire was administered in which participants were asked “Have you ever seen anybody for emotional or psychiatric problems?” and “Was there ever a time when you, or someone else, thought you should see someone because of the way you were feeling or acting?”. If they answered ‘yes’ to either of these questions, they were asked to complete the SCID (22). If they answered ‘no’ to both questions, they were assigned control status. Those who completed the SCID but did not meet the criteria for MDD or other major psychiatric disorder were also defined as controls. Individuals who declined to complete the screening questionnaire or SCID were not included in further analysis. Individuals were also asked about smoking and drinking status. Individuals were asked ‘Have you ever smoked tobacco?’, or ‘Have you ever had an alcoholic drink?’. They were requested to respond according to: (i) Yes, currently smoke/drink, (ii) Yes, but stopped within past 12 months, (iii) Yes, but stopped more than 12 months ago, (iv) No, never smoked/drank. BMI was calculated using height (cm) and weight (kg) measured by trained clinical staff.

#### Derivation of the epigenetic clock

Whole blood genomic DNA (500 ng) samples from 5,200 individuals were treated with sodium bisulphite using the EZ-96 DNA Methylation Kit (Zymo Research, Irvine, California), following the manufacturer’s instructions. DNA methylation was assessed using the Infinium MethylationEPIC BeadChip (Illumina Inc., San Diego, California), in accordance with the manufacturer’s protocol. The arrays were scanned using a HiScan scanner (Illumina Inc., San Diego, California) and initial inspection of array quality was carried out using Genome Studio v2011.1.

Additional quality control measures were implemented using the R packages ‘shinyMethyl’ (23, 24) and ‘wateRmelon’ (23). First, shinyMethyl was used to plot the log median intensity of the methylated signal against the log median intensity of the unmethylated signal for each array. Outliers from this QC plot were visually identified and excluded from further analysis. Methylation beta-values were then entered into the ‘pfilter’ function in wateRmelon which was used to exclude poor-performing samples and probes. Samples were excluded if ≥ 1% sites had a detection p-value of > 0.05. Probes were removed from the dataset if: (i) they had more than 5 samples with a beadcount of less than 3; or (ii) ≥ 0.5% samples had a detection p-value of > 0.05. Finally, shinyMethyl’s sex prediction plot was used to exclude samples whose predicted sex differed from their recorded sex. After QC, the dataset comprised beta-values for 860,926 methylation loci measured in 5,101 individuals.

The DNAm age measures were calculated using the coefficients reported in the Hannum and Horvath publications (10–12). Of the 71 CpGs in the Hannum clock, 65 were included in the QCd current dataset and 335 of the 353 Horvath clock CpGs. Correlations between the clocks with chronological age in a sub-sample of 2,586 individuals (selected to be unrelated and not share a nuclear environment; no couples and no pairs with pi-hat>0.05) are shown in Supplementary Figure 1, r=0.94 and r=0.91 for Hannum and Horvath clocks respectively. As an additional QC measure, we performed analysis excluding data where epigenetic age values fell out-with three standard deviations from the group mean. The current sample of individuals with QC’d epigenetic data and MDD status data therefore included n_Hannum_ =3,830 controls, n_Hannum_=1,217 MDD cases and n_Horvath_=3,833 controls, n_Horvath_=1,219 MDD cases.

Measures of 'epigenetic age acceleration' (EAA) were derived from both the Horvath and Hannum clocks, by regressing DNAm age on chronological age and taking the corresponding residuals as the EAA measure. A positive EAA value indicates that DNAm age is higher than chronological age (accelerated ageing), and negative EAA values represent younger biological age compared to chronological age.

#### Statistical modelling

All analyses were conducted in ASReml-R (www.vsni.co.uk/software/asreml, version 3.0). Associations were examined between EAA as the dependent variable and MDD status as the independent variable, controlling for sex using mixed linear model association analysis. Since GS:SFHS is a family-based study, family structure was fitted as a random effect by creating an inverse relationship matrix using pedigree kinship information to control for relatedness (25). These covariates (sex, relatedness), as well as measures of blood cell-type composition (including CD8T, CD4T, NK, B cells and granulocytes, estimated using minfi’s ‘estimateCellCounts’ function) and processing batch, were included in the basic model (13, 14).

Further covariates were also added to test sensitivity of the regression model to potential confounding lifestyle and health-related factors previously associated with DNAm (full models). These included self-reported smoking and alcohol intake/consumption (as described above). In addition, since there is a known relationship between obesity and accelerated epigenetic ageing (26), and between obesity and MDD (27), we also explored the contribution of body mass index (BMI, as a proxy for obesity) to the relationship between EAA and depression status.

Additional models to account for potential interaction effects of covariates (age and sex) on MDD status (e.g. modelling MDD*age, MDD*sex) are contained within supplementary materials (Supplementary Tables 1a.b).

Out of the three additional covariates (smoking, drinking status and BMI), BMI was the only one that demonstrated an attenuating effect on the relationship between EAA and MDD status. We therefore further examined the shared genetic aetiology between EAA, MDD and BMI. To test the potential overlap in polygenic architecture between EAA, MDD and BMI, and estimate the size and significance of any genetic correlation, we used cross-trait Linkage Disequilibrium Score regression (LD Score regression) (28, 29). We followed the protocol outlined by Bulik-Sullivan et al. (29), using data from GWAS analyses of EAA_Hannum_ (n=5047) and EAA_Horvath_ (n=5052) from the current sample (see supplemental materials), summary statistics from the recent GWAS of BMI from the GIANT consortium (30), and summary statistics from the largest available GWAS of MDD (N=453,779, of which 129,667 were MDD cases), carried out by the Psychiatric Genomics Consortium (31), which includes summary statistics from the personal genetics company 23andMe, Inc., (32) and UK Biobank (with Generation Scotland samples removed).

We additionally conducted a formal mediation analysis of the relationship between EAA and MDD, as mediated through BMI, using the ‘testMediation’ function in R using the Sobel method on an unrelated sample of individuals (controls n=1,301, MDD cases n=463). In each case EAA was entered as the independent variable, MDD status was entered as the dependent variable, and BMI was entered as the mediating variable. Covariates included were sex, cell counts and batch, along with smoking and drinking status. For completeness, we also tested for mediation effects of smoking and drinking status on the relationship between EAA and depression status, see supplementary materials.

For all analyses, all continuous measures were rescaled into zero mean and unitary standard deviation in order that effect sizes represent standardized scores.

## Results

### Demographics

Sample characteristics are presented in Table 1. There was no significant difference in chronological age between cases and controls (t=-0.2131, p=0.83). Individuals with MDD were more likely to be female (χ^2^ = 74.864, p < 2.2×10^−16^), more likely to be current smokers (χ^2^ = 30.251, p = 1.2×10^−6^), and less likely to be current drinkers (χ^2^ = 56.645, p = 3.06×10^−12^) than control individuals (see Table 1). Body mass index (BMI) was greater in MDD cases than controls (controlling for sex, β=0.1106, p=8.38×10^−4^), see Table 1.

**Table 1:** Sample Characteristics

| | Controls (N = 3,833) | | MDD cases (N =1,219) | | $\beta/\chi^2/t$<br>(p value) |
| --- | --- | --- | --- | --- | --- |
|  | Mean | Std Dev | Mean | Std Dev |  |
| **Demographic variables and Epigenetic Age** |  |  |  |  |  |
| Age | 48.10 | 14.50 | 48.02 | 12.08 | 0.2131 (0.83) |
| Gender* (F:M), %F | 2228:1605 | 58% | 878:341 | 72% | 74.86<br>($2.2 \times 10^{-16}$) |
| Hannum age | 45.53 | 10.5 | 45.46 | 8.70 | See table 2 |
| Hannum age accel | 0.0006 | 3.17 | -0.0019 | 2.97 | See table 2 |
| Horvath age* | 52.49 | 9.83 | 52.70 | 8.23 | See table 2 |
| Horvath age accel* | -0.0652 | 3.53 | 0.2050 | 3.40 | See table 2 |
| **Symptoms, lifestyle and health-related variables** |  |  |  |  |  |
| Smoking status%*# | 17.7; 3.2; 25.6; 53.5 | - | 22.8; 3.2; 29.3; 44.7 | - | 30.251<br>($1.20 \times 10^{-6}$) |
| Drinking status%*# | 91.5; 1.6; 4.0; 2.9 | - | 86.1; 2.2; 9.5; 2.2 | - | 56.645<br>($3.06 \times 10^{-12}$) |
| BMI* | 26.88 | 5.16 | 27.52 | 5.96 | 0.1106<br>($8.38 \times 10^{-4}$) |

\*Significantly different ( $p < 0.05$ ) between groups, see text.

### MDD Status and Epigenetic Age Acceleration (EAA): Basic Models

Epigenetic age and epigenetic age acceleration measures are presented in Table 1 for both clock-based methods for MDD cases and controls. No group differences were found for EAA_Hannum_ in any of the models (Table 2). The EAA_Horvath_ however indicated significantly increased epigenetic age acceleration in depressed cases versus controls, see Figure 1, Table 2, with an overall mean positive value in cases (accelerated ageing, equivalent to 0.20 years) and a mean negative value (decelerated ageing, equivalent to 0.07 years) in controls. For the EAA_Horvath_ analysis controlling for relatedness and sex, the effect size was β=0.1103, p<0.001. In the analysis controlling for relatedness, sex, as well as processing batch and cell counts, the effect size was β=0.0804, p=0.012.

**Figure 1:**
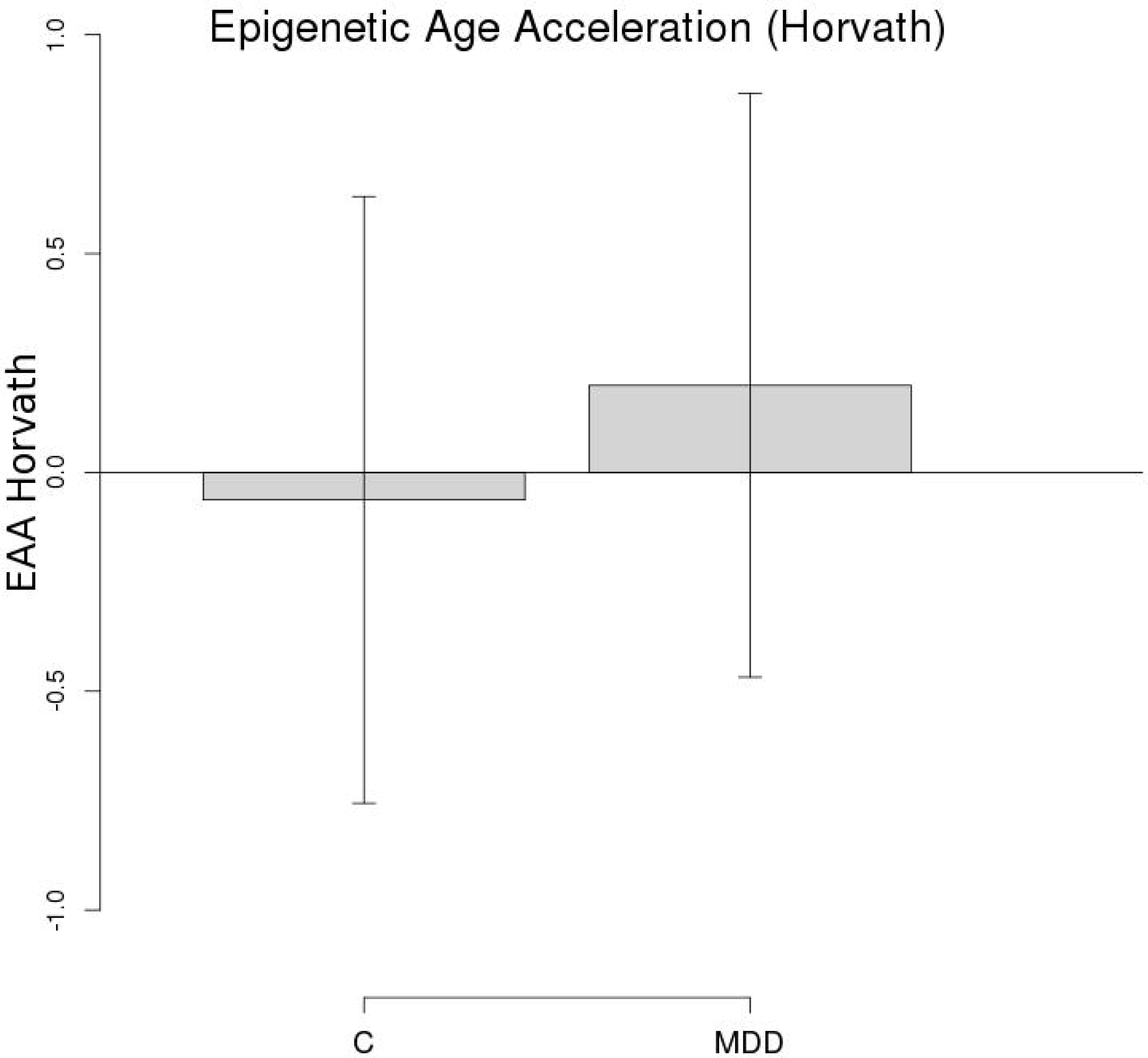
Boxplots of EAA (Horvath) according to MDD status. Boxplot of Epigenetic Age Acceleration according to the Horvath clock according to case-control status with mean and 95 % CIs

**Table 2:** Epigenetic Age Acceleration in MDD Cases versus Controls

|  | MDD effects |  |  |  |
| --- | --- | --- | --- | --- |
| EAA Horvath clock (basic model): | β |  | P value |  |
| Controlling for relatedness, sex | 0.1103 |  | 8.41x10<sup>-4</sup> |  |
| Controlling for above plus cell counts and batch | 0.0804 |  | 0.012 |  |
| EAA Horvath clock (full model):<br>incremental inclusion of covariates |  |  |  |  |
| Controlling for above plus smoking | 0.0811 |  | 0.013 |  |
| Controlling for above plus drinking | 0.0765 |  | 0.020 |  |
| Controlling for above plus BMI | 0.0647 |  | 0.049 |  |
| EAA Horvath clock (full model: separate inclusion of covariates) | MDD effects |  | Covariate effects |  |
|  | β | P value | β | P value |
| Controlling for relatedness, sex, cell counts and batch | 0.0804 | 0.012 | - | - |
| Controlling for relatedness, sex, cell counts, batch: & smoking only | 0.0811 | 0.013 | -0.0250 | 0.038 |
| Controlling for relatedness, sex, cell counts, batch: drinking only | 0.0834 | 0.011 | -0.0008 | 0.967 |
| Controlling for relatedness, sex, cell counts, batch: & BMI only | 0.0690 | 0.031 | -0.0900 | 7.35x10<sup>-11</sup> |

|  | MDD effects |  |
| --- | --- | --- |
| EAA Hannum clock (basic model): | β | P value |
| Controlling for relatedness, sex | 0.0456 | 0.165 |
| Controlling for above plus cell counts and batch | 0.0430 | 0.158 |
| EAA Hannum clock (full model):<br>incremental inclusion of covariates |  |  |
| Controlling for above plus smoking | 0.0351 | 0.255 |

*Accelerated epigenetic ageing in depression*
| Controlling for above plus drinking | 0.0308 |  | 0.322 |  |
| --- | --- | --- | --- | --- |
| Controlling for above plus BMI | 0.0208 |  | 0.503 |  |
| **EAA Hannum clock (full model): separate inclusion of covariates** | **MDD effects** |  | **Covariate effects** |  |
|  | **β** | **P value** | **β** | **P value** |
| Controlling for relatedness, sex, cell counts and batch | 0.0430 | 0.158 | - | - |
| Controlling for relatedness, sex, cell counts, batch: & **smoking only** | 0.0351 | 0.255 | -0.0759 | **6.96x10<sup>-11</sup>** |
| Controlling for relatedness, sex, cell counts, batch: **drinking only** | 0.0454 | 0.143 | -0.0295 | 0.131 |
| Controlling for relatedness, sex, cell counts, batch: & **BMI only** | 0.0350 | 0.251 | -0.0763 | **6.24x10<sup>-9</sup>** |

Figure 2 demonstrates a scatterplot of EAA_Horvath_ versus chronological age coded by MDD status, which indicated a greater difference in EAA_Horvath_ in MDD individuals versus controls at the younger age range of the cohort.

**Figure 2:**
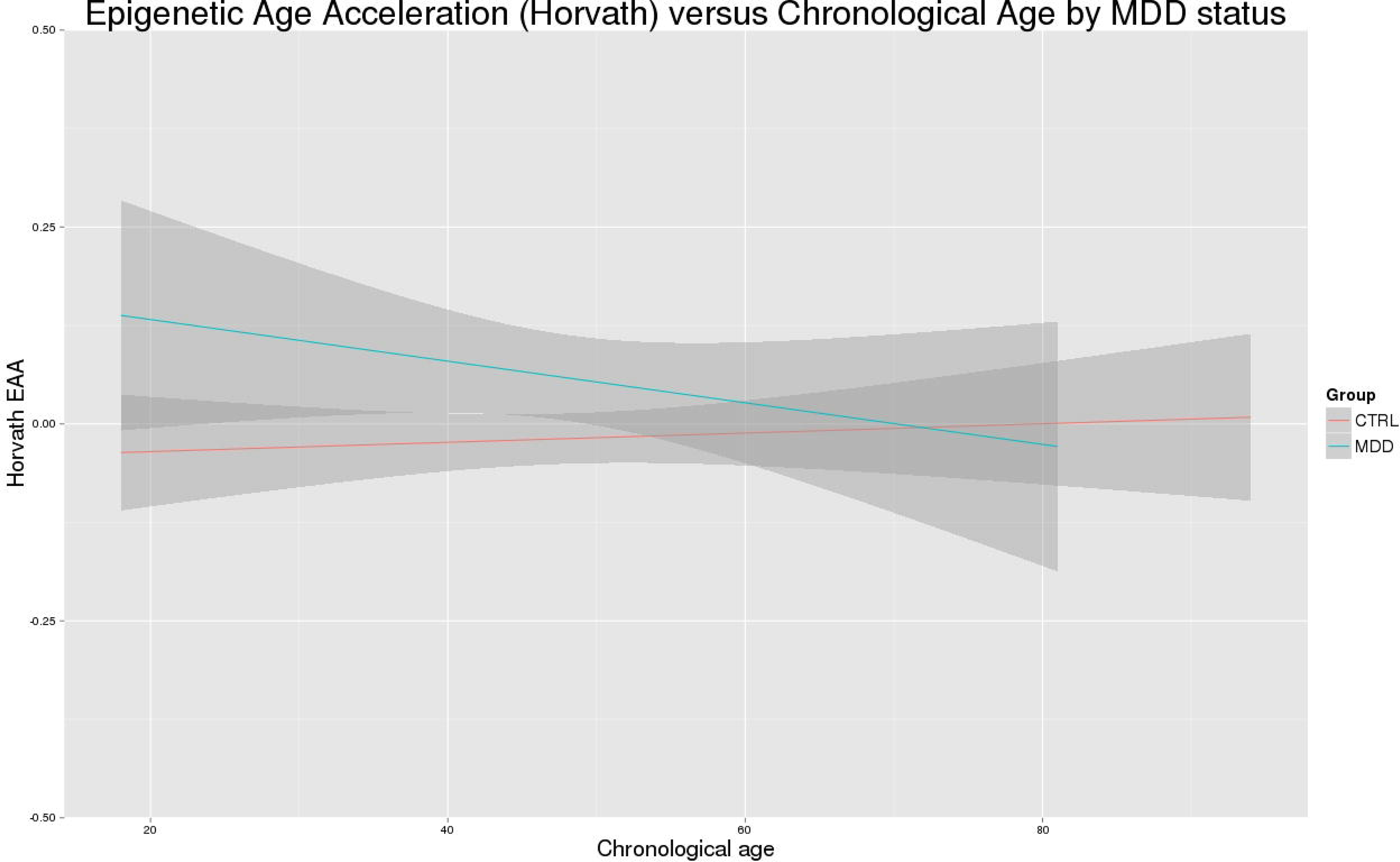
Scatterplots of EAA (Horvath) by chronological age, coded by depression status.

### MDD Status and Epigenetic Age Acceleration (EAA): Full Models

To determine whether observed increases in EAA_Horvath_ in MDD were due to potential confounding lifestyle and health-related factors, the previous statistical models were run including smoking status as an additional covariate. Here MDD status remained significantly associated with increased EAA_Horvath_ (β=0.0811, p=0.013). The next sensitivity analysis included all above covariates plus smoking and drinking status, and again MDD status remained significantly associated with increased EAA_Horvath_ (β=0.0765, p=0.020). With the incremental inclusion of BMI in the model the results were also still significant (β=0.0647, p=0.049).

Table 2 also demonstrates the effect of these additional covariates added separately to the model. Only smoking (β=-0.0256, p=0.039) and BMI (β=0.0886, p=2.22×10^−10^) demonstrated a significant effect in these models. However, only BMI demonstrated an attenuation of the effect size in relation to MDD status (from β=0.0804 to β=0.0690 with the addition of BMI, β=0.0804 to β=0.0811 with the addition of smoking, and β=0.0804 to β=0.0834 with the addition of drinking status).

### MDD * age/sex interaction effects

The inclusion of the age*MDD and sex*MDD interaction terms as covariates are presented in Supplementary Table 1a. The addition of these covariates either singly or together in the same model did not alter the main pattern of results described above for effects of MDD status (increased EAA in MDD cases), with the exception of the full models incrementally including all covariates plus BMI where results fell to just below significant (β=0.0644, p=0.051). Notably, the MDD*age interaction term was itself significant in these full models (β=-0.0723 to β=-0.0952, p<=0.05), as reflected in the divergence of the groups at the younger age range of the cohort, depicted in Figure 2. MDD*sex interactions were not significant.

### Genetic correlation analysis: EAA, MDD and BMI

Genetic correlation analysis indicated there was a significant shared genetic architecture between EAA_Horvath_ and BMI (r_G_=0.20, p=0.031), and between MDD and BMI (r_G_=0.10, p=9.86×10^−6^, see Table 3, Figure 3), but not between EAA_Horvath_ and MDD (r_G_=0.14, p=0.125). The genetic correlation between EAA_Hannum_ and EAA_Horvath_ also indicated shared architecture between the two measures of accelerated ageing (r_G_=0.68, p=0.035), though there were no significant genetic correlations between EAA_Hannum_ and MDD or BMI.

**Table 3:** Results of cross-trait LD Score regression analysis

| Trait 1 | Trait 2 | Trait 1 heritability | Trait 2 heritability | Genetic correlation | P-value |
| --- | --- | --- | --- | --- | --- |
| EAA<sub>Horvath</sub> | MDD | 0.2098 (0.1002) | 0.0881 (0.0039) | 0.1387 (0.0913) | 0.125 |
| EAA<sub>Horvath</sub> | BMI | 0.2104 (0.1075) | 0.1391 (0.0079) | 0.1962 (0.0907) | **0.031** |
| MDD | BMI | 0.0892 (0.0036) | 0.1391 (0.0082) | 0.1007 (0.0228) | **9.86x10<sup>-6</sup>** |
| EAA<sub>Horvath</sub> | EAA<sub>Hannum</sub> | 0.1779 (0.0928) | 0.1147 (0.0889) | 0.6807 (0.3234) | **0.035** |
| EAA<sub>Hannum</sub> | MDD | 0.0972 (0.0862) | 0.0881 (0.0039) | 0.1381 (0.1603) | 0.326 |
| EAA<sub>Hannum</sub> | BMI | 0.0886 (0.0846) | 0.1390 (0.0079) | 0.2287 (0.1568) | 0.145 |

Note: MDD data from PGC BMI data from GIANT, see methods for further details. Heritabilities reported are on the observed scale for continuous traits (age acceleration and BMI) and on the liability scale for MDD. Significant p-values ( $p < 0.05$ ) are highlighted in bold.

**Figure 3:**
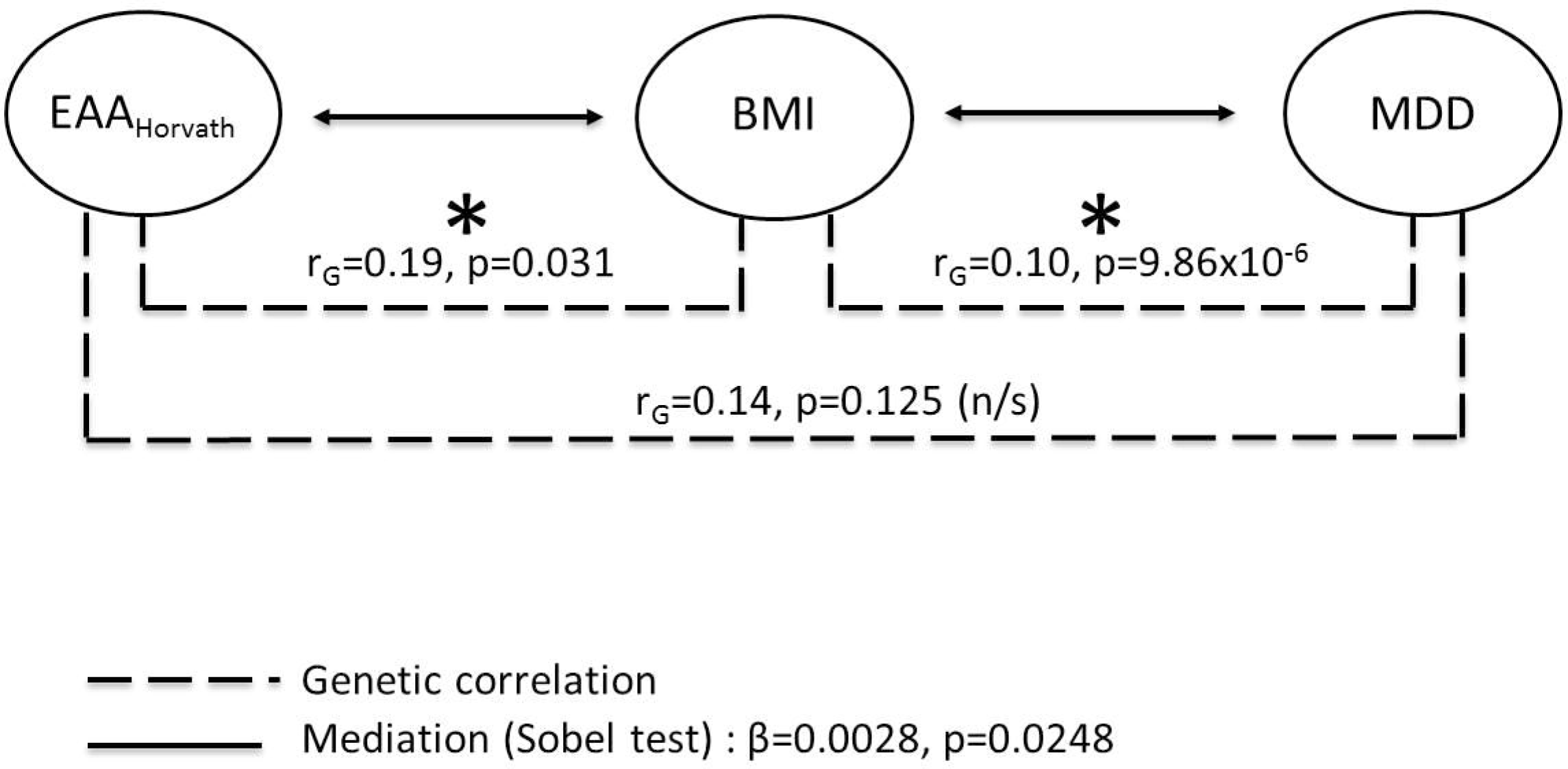
Mediation model and genetic correlation summary between EAA (Horvath), BMI and MDD.

### Mediation analysis: EAA, MDD and BMI

Tests of formal mediation suggested that BMI partially mediated the relationship between EAA and depression status. Specifically, the Sobel test showed a significant mediating effect of BMI between EAA_Horvath_ and depression status (β=0.0028, p=0.025, see Figure 3 and Supplementary Table 2). The proportion of the effect mediated by BMI (indirect effect/total effect) was 12.79%. Mediating effects of smoking and drinking did not significantly mediate this relationship (β=0.0006, p=0.291, 4.11%, β= <0.0001, p=0.251, <0.01% respectively).

## Discussion

To our knowledge this is the largest study to date (>5,000 individuals) of epigenetic ageing in depressed cases versus controls. As hypothesised, we demonstrate that MDD is associated with a significantly higher EAA_Horvath_, contributing to the increasing body of evidence indicating accelerated biological ageing as a core feature of the disorder. The analysis further indicated that the observed epigenetic age acceleration effects in MDD were not due to confounding effects related to processing batch or cell counts, and were not attributable to smoking or drinking status. This difference between cases and controls was small (equating to 0.20 years), which may in part relate to the extensive heterogeneity associated with MDD (33). In contrast to our expectation that differences between MDD cases and controls would increase with age, we found larger differences in EAA towards the younger end of the cohort’s age range (see Figure 2).

Numerous lines of evidence have previously led to suggestions of accelerated ageing effects in MDD. These include observations that depressed individuals demonstrate an increased risk of ageing-related diseases and reduced life expectancy (3), overlapping biological pathways involved in both MDD and biological ageing (5), and evidence of excessive cellular ageing (2, 4, 34). Ageing and ageing-related diseases are also notably associated with changes in DNA methylation, leading to the development of DNA methylation-based predictors of biological ageing (10–12). It was previously unknown however, whether accelerated ageing effects at the level of methylation using such predictors would be seen in MDD, and how these may be mediated by other factors, for example BMI. There are known robust reciprocal links between obesity and depression (26), and the role of increased BMI and shortened lifespan has also been extensively studied (35). More recently, studies have also reported that increases in BMI are associated with accelerated epigenetic ageing, although mechanisms are currently unknown, as are the links with comorbid diseases such as MDD (26).

The current results support the idea that there is increased epigenetic age acceleration in depressed cases versus controls. We found evidence that these relationships were partially mediated by higher BMI (~13%), but not with smoking or drinking status. Future research probing the potential contributions of environmental and lifestyle influences on psychological well-being in the general population could highlight the importance of risk-related and protective factors of wide applicability.

Notably our findings between the two different DNAm age predictors differed in their sensitivity with regards the categorical depression diagnosis. It is notable however that there are differences between these two DNAm age methods that may contribute to these inconsistencies. The Hannum predictor is derived from whole blood DNA from a single cohort of individuals using 71 CpGs. The Horvath model was however derived from multiple tissue types using data from multiple independent studies and uses 353 CpGs. Further, each method involves almost entirely non-overlapping CpG sites. Although both methods perform similarly in their prediction of mortality (14, 36), previous studies have also reported differences in their associations between other traits of interest, for example exposure to environmental particulate matter (37), with neural integrity in post-traumatic stress disorder (36), and with longevity (38). These studies, together with the current findings, therefore suggest that the two DNAm age biomarkers of biological ageing may reflect subtly different aspects of the ageing process.

The major strength of our study is the large sample size, with DNA methylation data available on more than 5,000 individuals, plus the availability of detailed phenotyping in our cohort. As sample sizes for GWAS of EAA increase, future work could include genetic analyses to inform causation models, such as Mendelian Randomisation. The lack of access to brain tissue is a frequently cited limitation of blood-based DNA methylation studies. Whilst we are unable to make strong inferences specifically about effects on the brain, other studies have indicated its suitability as a relevant for biomarker for such research purposes (11, 39). Secondly, this study is cross-sectional in nature therefore it is not possible to infer causal directionality between factors. Lastly, although we corrected for several confounders that might influence DNA methylation, such as sex, age*MDD, sex*MDD, smoking, drinking, and BMI, we cannot exclude the possibility that other factors not captured by our methods may have confounded the observed relationships.

In conclusion, our results indicate a significant increase in epigenetic age acceleration in depressed cases versus controls lending further support to the hypothesis of accelerated biological ageing in MDD. Since this effect was greatest for the younger individuals it is possible that this may also represent altered maturation. Our findings extend previous findings to suggest that the positive relationship between accelerated methylation age and depression may be partially mediated through increased BMI. Future research would be needed to determine the direction of causality, and whether these associations can be reversed or mitigated by treatment or lifestyle interventions.

## Acknowledgements

This study is supported by a Wellcome Trust Strategic Award “Stratifying Resilience and Depression Longitudinally” (STRADL) (Reference 104036/Z/14/Z) and by the Sackler Foundation. Generation Scotland received core support from the Chief Scientist Office of the Scottish Government Health Directorates [CZD/16/6] and the Scottish Funding Council [HR03006]. Genotyping of the GS:SFHS samples was carried out by the Genetics Core Laboratory at the Wellcome Trust Clinical Research Facility, Edinburgh, Scotland and was funded by the Medical Research Council UK and the Wellcome Trust (Wellcome Trust Strategic Award (STRADL; Reference as above). HCW is supported by a JMAS SIM fellowship from the Royal College of Physicians of Edinburgh and by an ESAT College Fellowship from the University of Edinburgh. SRC is supported by a Medical Research Council (MRC) grant (MR/M013111/1). Part of the work was undertaken in The University of Edinburgh Centre for Cognitive Ageing and Cognitive Epidemiology (CCACE), part of the cross council Lifelong Health and Wellbeing Initiative (MR/K026992/1); funding from the Biotechnology and Biological Sciences Research Council (BBSRC) and MRC is gratefully acknowledged. Age UK (The Disconnected Mind project) also provided support for the work undertaken at CCACE. We would also like to thank the research participants and employees of 23andMe for making this work possible. We thank the following members of the 23andMe Research Team: Michelle Agee, Babak Alipanahi, Adam Auton, Robert K. Bell, Katarzyna Bryc, Sarah L. Elson, Pierre Fontanillas, Nicholas A. Furlotte, David A. Hinds, Karen E. Huber, Aaron Kleinman, Nadia K. Litterman, Jennifer C. McCreight, Matthew H. McIntyre, Joanna L. Mountain, Elizabeth S. Noblin, Carrie A.M. Northover, Steven J. Pitts, J. Fah Sathirapongsasuti, Olga V. Sazonova, Janie F. Shelton, Suyash Shringarpure, Chao Tian, Joyce Y. Tung, Vladimir Vacic, and Catherine H. Wilson.

